# Eco-evolutionary agriculture: a study in crop rotations

**DOI:** 10.1101/402313

**Authors:** Maria Bargués-Ribera, Chaitanya S. Gokhale

**Affiliations:** Research Group for Theoretical Models of Eco-evolutionary Dynamics, Department of Evolutionary Theory, Max Planck Institute for Evolutionary Biology, August-Thienemann-Straβe 2, 24306 Plön, Germany.

**Keywords:** agriculture, crop rotations, eco-evolutionary dynamics, plant-pathogen co-evolution

## Abstract

Since its origins, thousands of years ago, agriculture has been challenged by the presence of evolving plant pathogens. In response, current practices have started relying on computational tools to design efficient prospective planning, but further efforts for multi-criteria assessment are needed. Here, we present a methodology for developing cultivation strategies optimal for control or eradication of pathogens. This approach can integrate both, traditionally used criteria in crop rotations and the analysis of host-pathogen coevolution systems where hosts are artificially selected. Our analysis shows that prospective planning can maximise cash yield in the long run by investing consecutively in soil quality during initial seasons. Importantly, rational application of crop rotation patterns can minimise yield loss in infected fields, despite the evolution of pathogen virulence. Our results provide strategies for optimal resource investment for increased food production and lead to further insights into minimisation of pesticide use in a society demanding efficient agriculture.

## Introduction

Around ten thousand years ago, changes in climate conditions led to the emergence of agricultural practices in human hunter-gatherer communities around the globe [1]. This process of domestication – or artificial selection – has been refined along centuries through trial and error combined with experience, increasing the quantity and quality of the product. In the case of plant agriculture, the presence of pests has been a substantial threat to farmers [2]. The first farmers already tried to overcome the pest problem by employing field rotations, i.e., shifting cultivation techniques [3, 4]. As human population continues to multiply, current agriculture practices need to address a two-fold problem of the dearth of enough food supply and plant pathogens. Techniques such as slash-and-burn, pesticides and fertilisers are used for increasing yield as well as dealing with pests, but do not contribute to agricultural sustainability [5]. Thus, current research needs to focus on developing cropping techniques which increase yield and minimise the environmental impact [6]. Farmers have now started relying on data-based computational tools to design agriculture strategies. Among others, the computational tools involve decision support models for choosing optimal cropping plans and crop rotation decisions [7, 8]. The models guide allocating crops depending on their characteristics – botanical family, market demand, or soil demands –, considering spatial distribution and temporal successions. However, these models are presented as static decisions and the optimisation procedure often lacks multicriteria assessment [9] one such being plant pathogens.

In evolutionary biology, models on host-pathogen coevolution have contributed to understanding the relationship between some pathogens and their host crops, for example, regarding the specificity of the interaction [10, 11]. Plant-pathogen coevolution is a natural selection process happening in tandem with artificial selection of domestication, and the interface between them is worth studying. Recently, authors have highlighted the use of plant-pathogen coevolution in formulating disease management strategies and avoiding the increase of infectivity in pathogens [12, 13, 14, 15, 16]. When observing evolutionary dynamics, a change in the crop type acts as a per-turbation leading to frequent selective sweep-like dynamics. Tracking the frequency and speed of such sweeps would be useful in detecting periods of lower fitness and reduced population size; in which the pathogen could be pushed to extinction [17]. Thus, including plant-pathogen coevolution – a natural selection process – in the study of crop rotations would be a new approach for tackling agricultural problems.

Here, we aim to design a cultivating strategy optimal for pathogen eradication, integrating criteria traditionally used on crop rotations, and host and pathogen coevolution. We have developed an optimisation model for crop rotation patterns which appoints soil quality and cash yield value as variables of interest. When infection occurs, ecological dynamics play out in the short term. Host-pathogen competitive dynamics predicts yield loss depending on crop resistance, as well as changes in pathogen load. Evolution of host resistance and pathogen virulence is considered by mutation of the pathogen into strains which are able to infect the host more efficiently. While some crop rotation patterns are optimal in pathogen-free scenarios, different patterns are shown to be robust to infections, despite minor damages or sub-optimal yield output. Improvement strategies are proposed by (i) introducing engineered resistant variants of the original crops and analysing the corresponding pathogen adaptation following previous models [18]; and (ii) proposing new rotation sequences which show better performance in infection conditions.

## Model and Results

Plants have a variety of responses to pest infestations such as susceptibility, tolerance and resistance. In agriculture, farmers have used this variability for thousands of years to control the spread of pathogens over the field. This simple yet powerful concept is formalised below, using a theoretical model which optimises rotation patterns and predicts infection dynamics.

### 2.1 Crop rotation patterns

To establish a basic model of rotation patterns, we focus on sequential combinations of cash crops and cover crops. Cash crops are those which provide a product to be commercialised – e.g. maize –, whereas cover crops improve soil quality of the field but provide no direct substantial cash yield – e.g. clover. Because more soil quality provides more cash yield, both crop types are needed for successful farming: the aim of the basic model is to study which temporal patterns of cash and cover crops maximise the farmer’s benefit. For that, we have been inspired by a previous report assessing the optimal length of clover period, compared to maize, in a 9-year field experiment [19]. We work with pattern sequences of length *L* = 10, in contrast with common short rotations of *L* ≈ 3, aiming to acquaint long-term patterns with increased optimisation.

The more cash yield a particular sequence provides the farmer, the more likely it is to be used repeatedly. This concept can be connected to the idea of differential survival. Better yielding sequences survive and reproduce i.e. are used again. Thus, we can connect the notion of fitness to the optimality of a sequence [20]. Here we attribute each of our rotation sequences a fitness value *F*, depending on their ability to maximise cash yield. In principle, the fitness function could be modified to optimise both final soil quality and cash yield, as discussed later on.

Each element in a rotation sequence corresponds to a harvesting season, modelled as a discrete time-step. During each time-step, there is a change in soil quality *q*_*t*_ and cash yield *y*_*t*_, which varies depending on the host type *h*_*i*_, *i* = {1, 2}. At the end of the sequence, *F* is set as the final cash yield (Fig. 1). Soil quality and cash yield per time step are crucial in obtaining the fitness value, taking the following form:

**Figure 1:**
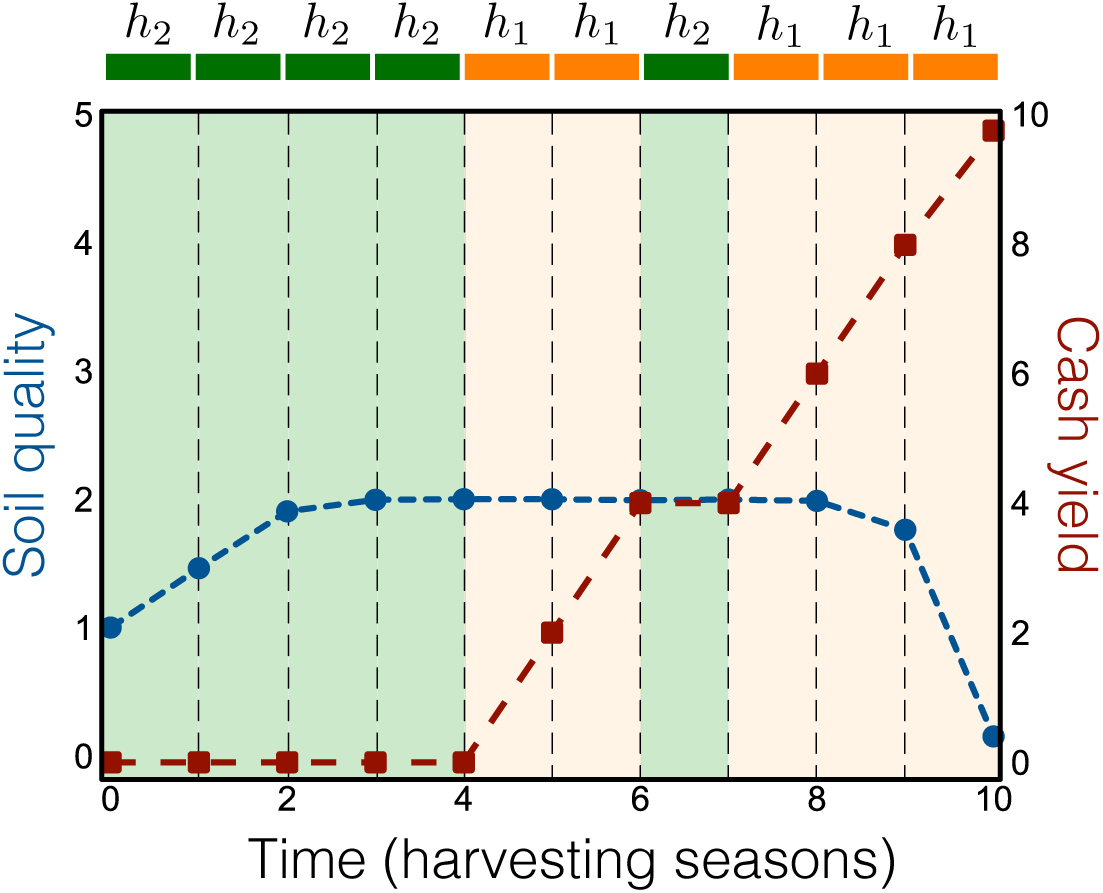
Soil quality and cash yield values of a rotation sequence along time. Each time step corresponds to a harvesting season. Dots indicate discrete values for soil quality (circles) and cash (squares). Season crop type is indicated in orange for cash crops *h*_1_, and green for cover crops *h*_2_. Fitness *F* is equivalent to the cash yield value at *t* = *L* (here *t* = 10).

- Soil quality (*q*_*t*_): Soil quality decreases following a logistic decay curve for *n*_1_ consecutive *h*_1_ cash crops, and increases with logistic growth for *n*_2_ consecutive *h*_2_ cover crops. The parameter *β*_*i*_ regulates the intensity and direction of the soil quality change given crop type *i*: we set *β*_1_ = *-*1.5 for soil quality decrease by *h*_1_, and *β*_2_ = 1 for soil quality increase by *h*_2_, considering that it is more difficult to improve soil quality than to decrease it (see Supplementary Material). We assume that the soil quality cannot increase indefinitely, reaching a saturation value – or carrying capacity – of *K*. We choose *K* = 2, for which approximately *n*_2_ = 4 harvesting seasons are needed to reach it with *β*_2_ = 1, similar to experimental work [19].

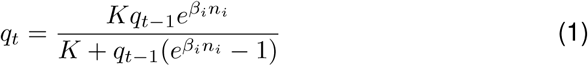
- Cash yield (*y*_*t*_): Cash yield increases in proportion to the soil quality of the previous time-step *q*_*t-*1_, regulated by the effective crop ratio *δ*_*i*_, and the crop yield contribution *γ*_*i*_. For cash crops we set *γ*_1_ = 1, considering cash yield increases as much as the soil quality, in a 1:1 ratio. For cover crops, there is no cash yield, being *γ*_2_ = 0. Overall cash yield is accumulated along the rotation sequence time-steps.

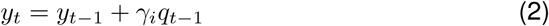
- Fitness (*F*): Given that we consider fitness to be the total economical benefit that a sequence provides, we set it equivalent to the total accumulated cash yield (i.e. *y*_*L*_).

Fitness of each sequence in a population of sequences of length *L* = 10 is computed according to the above-defined functions and analysed.

Results show that fitness distribution of the population of rotation sequences of *L* = 10 (*N* = 1024) has a non-zero (positive) skew: the proportion of sequences with high fitness values is small, indicating it is crucial to know the patterns underlying them (Fig. 2a). We have analysed the selection of sequences with a fitness value greater than two standard deviations from the mean, with the sequence population being reduced to *N* = 16. From this selection, we can observe the following patterns: (a) all sequences in the selection have an equal proportion of cover and cash crops; (b) all sequences in the selection end with a minimum of two cash crops; (c) most sequences start with cover crops, and the ones that start with cash, they set consecutive cover crops in the next seasons (Fig. 2b, 2c).

**Figure 2:**
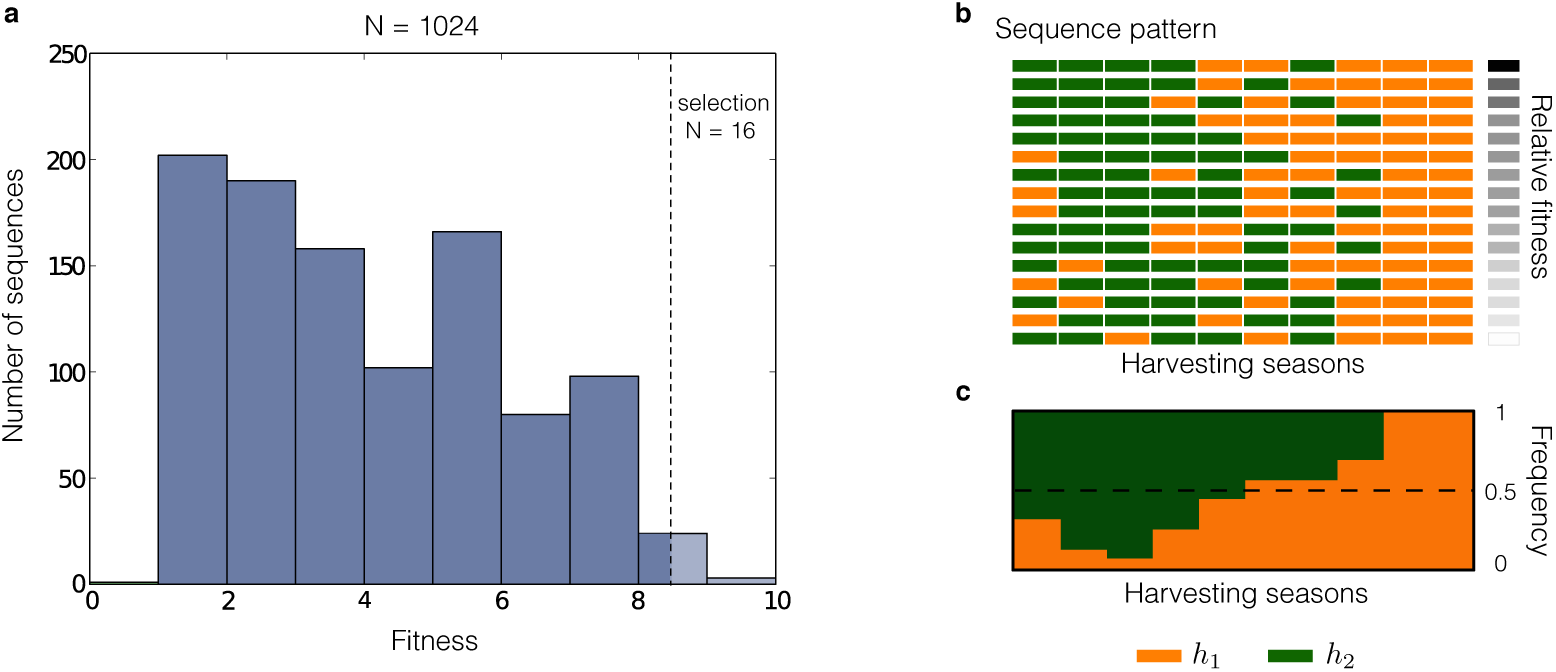
Fitness distribution of the population of sequences of *L* = 10 and its optimal patterns. a) Fitness distribution of population of study (*N* = 1024). Dashed line indicates two standard deviation over the mean threshold for *F.* b) Crop rotation patterns of selected sequences over the threshold (*N* = 16). Gray-scaled squares on the right indicate fitness value relative to the selection. c) Frequency of cash *h*_1_ and cover *h*_2_ crops among the selection of sequences in each harvesting season.

### 2.2 Patterns under infection: eco-evolutionary dynamics

In natural settings, the process of coevolution between the host plant and its pathogen can lead to the cyclic evolution of host resistance and parasite virulence, maintaining genetic diversity [14]. In agriculture, humans are the selecting agent: they decide which host grows in the next generation. While being economically important, the selected crop can be particularly vulnerable to pathogens which it has not coevolved with. Moreover, there are only a few major agricultural cash crops; resulting in less genetic diversity in host crops and more disease susceptibility [21]. In this section, we see how the introduction of a pathogen into our system modifies the fitness of the rotation sequences. We start with a simple infection scenario in which a pathogen *p* can infect cash crops *h*_1_, but not the cover crops *h*_2_. As an example, the fungi *Fusarium graminearum* [22] is one such pathogen which infects cash crops as maize or wheat, but does not infect cover crops such as clover.

**Ecological dynamics.** To include host-pathogen ecological dynamics, we adapt the Lotka-Volterra competitive equations, based on [23]. Within a season, time is continuous and dynamics are described by a system of ordinary differential equations, corresponding to *m* host types and *n* pathogens (Eq.(3)).

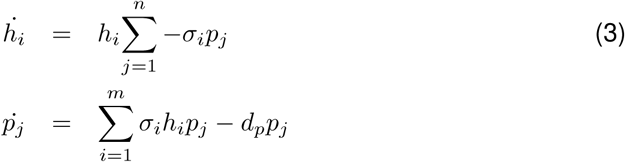

While the equations are for multiple hosts and pathogens, we start with two hosts and a single pathogen (*m* = 2 and *n* = 1). Here *h*_*i*_ is the population density of crop *i* and *p*_*j*_ is the pathogen density. The pathogen infectivity is set by *σ*_*i*_ (*σ*_1_ = 0.04 for the cash susceptible crop, *σ*_2_ = 0 for the cover resistant crop), and *d*_*p*_ is the death rate of the pathogen (*d*_*p*_ = 0.5). Due to the artificial setting of agriculture, we consider that without external perturbations and independently of the pathogen interaction, there is no birth nor death in host population during the season.

Between seasons, however, the system changes in discrete time-steps. At the end of each season, the crop is harvested, converted to yield and new crops are planted as per the rotation schedule. Due to the harvest, the pathogen population is disturbed. The pathogens are partially eliminated (*∊*) when the crop is harvested but some survive in the soil or in residues of the infected crops. We set *∊* = 0.5, considering half of the pathogen population stays in the field. Overall, the model evolves following continuous time within seasons, and discrete jumps between seasons, being an example of a hybrid dynamical system [24] (Fig. 3).

**Figure 3:**
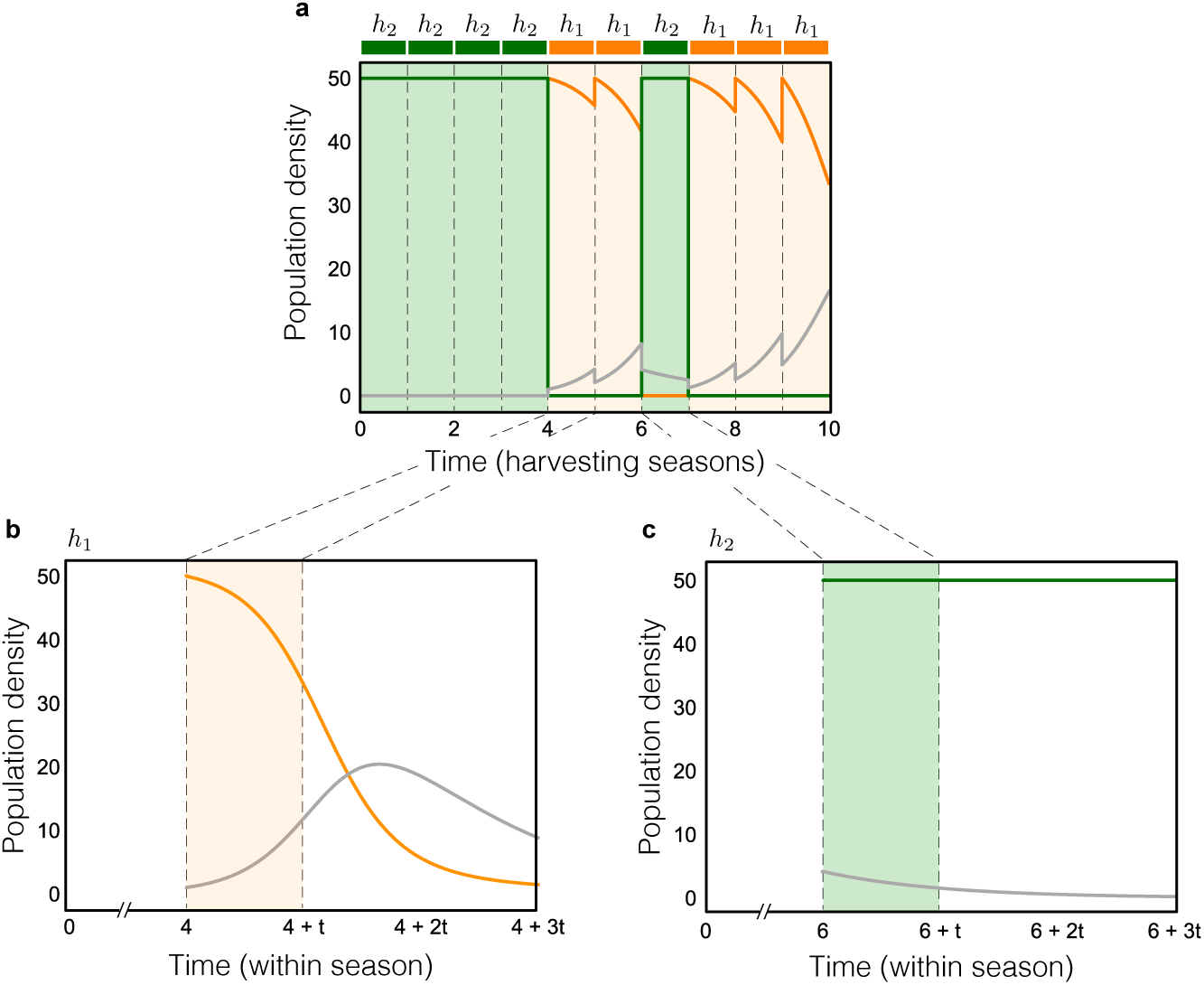
Host-pathogen ecological dynamics, between and within seasons. a) Dynamics between seasons, following a hybrid system. After each harvesting season, initial host density *H* = 50 is reinitialised and pathogen is reduced according to parameter *∊* = 0.5. b) Dynamics within a season, when there is a susceptible cash crop *h*_1_. Host density decreases due to the presence of the pathogen, while the pathogen load increases as long as there are enough crops to infect. c) Dynamics within a season, when there is a resistant cover crop *h*_2_. The resistant cover crop maintains its output throughout the season and remains unaffected by the presence of the pathogen; however, the pathogen dies out to to the lack of a suitable host to grow on.

**Eco-evolutionary Dynamics.** Substantial evolutionary changes can happen on ecological time-scales [25, 26, 27]; thus, we need to consider evolution in the hostpathogen interaction. In our case, we study the dynamics when there is evolution of pathogen virulence, and incorporate them in the already developed ecological dynamics. In literature, virulence refers to the pathogen capacity to establish an infection or the consequences for the host to be infected [28]. Here, by evolution of virulence, we refer to the change in the propensity of a pathogen to infect the host. Within a season, the pathogen reproduces, generates variation, and some of these variants may carry mutations that provide more virulence. To incorporate evolution in the ecological dynamics, we modify the previous equations (3) allowing the pathogen to mutate into strains which can exploit the cash host more efficiently, Eq. (4). The evolved pathogen cannot evolve to infect the cover crop since cash and cover are assumed to be phylogenetically distant.

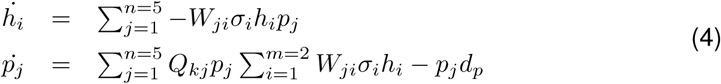

The new equations have two important elements: the transition matrix *Q*_*kj*_ and the fitness matrix *W*_*ji*_. The transition matrix *Q*_*kj*_ corresponds to the probabilities of the pathogen to mutate into other strains. We define four possible mutants, separated by a genetic distance *d* from the original strain. Mutation can happen between strains which are one mutational step away (*d* = 1) with a transition probability *µ* = 0.1.

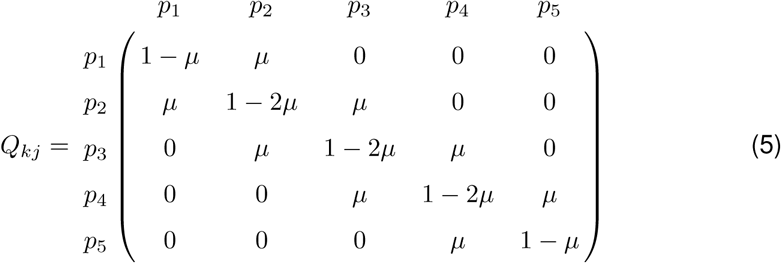

For the fitness matrix *W*_*ji*_, we set the fitness of the original pathogen *p*_1_ to *w*_11_ = 1 when infecting *h*_1_; and to *w*_12_ = 0 when infecting *h*_2_. Each mutant increases the fitness proportional to the distance with respect to *p*_1_, so *w*_*j*1_ = *w*_11_ + 0.25*d*, with *w*_51_ = 2 being the maximum fitness. Infecting *h*_2_ does not provide any fitness benefit, with *w*_*j*2_ = 0. The fitness matrix, when multiplied by the parameter *σ*, shows the increase in virulence in each mutated strain of the pathogen. Overall, eco-evolutionary dynamics results in elevated yield loss as compared to the solely ecological dynamics (Fig. 4).

**Figure 4:**
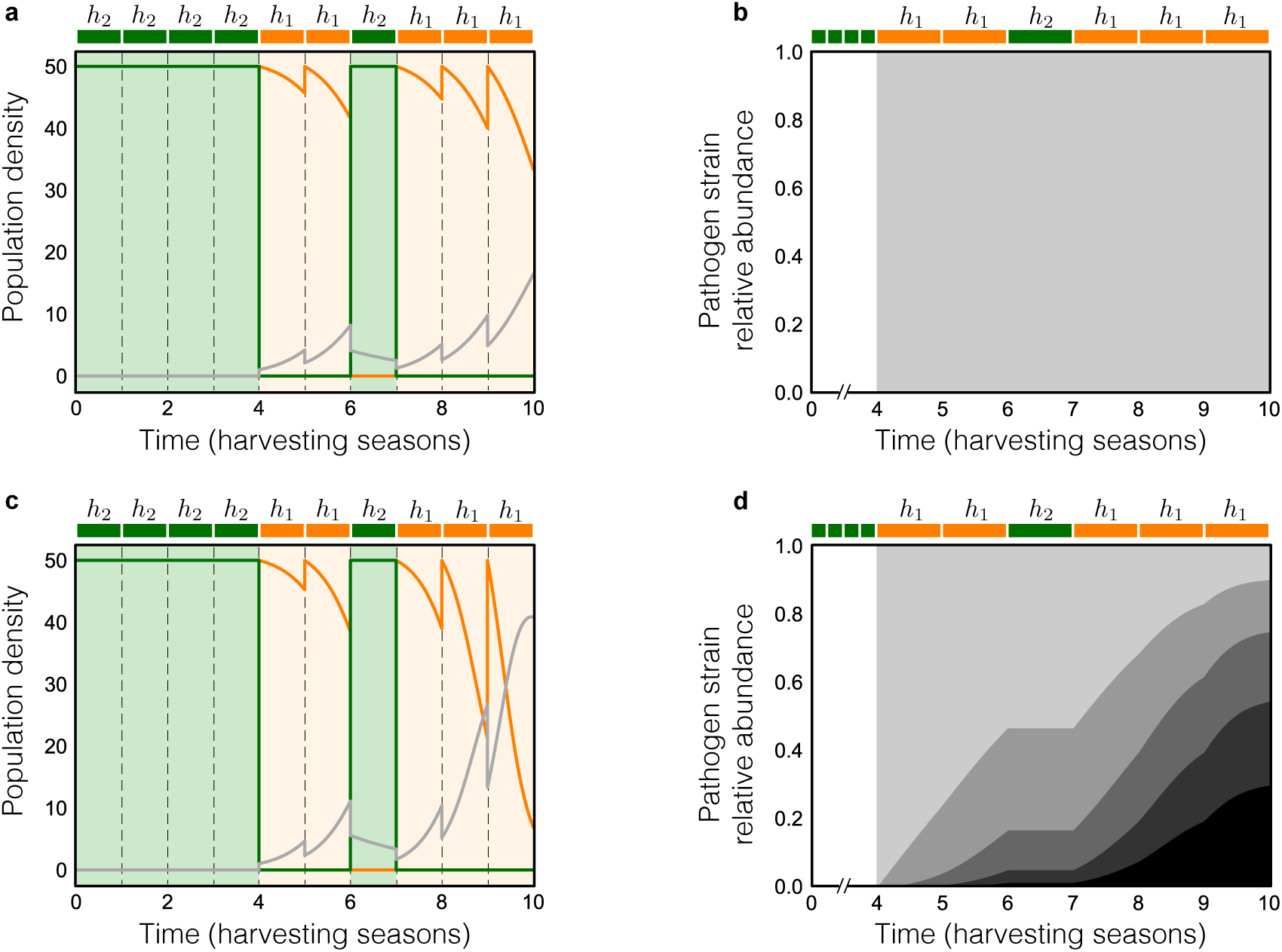
Ecological vs. eco-evolutionary dynamics of host-pathogen interaction. a) Ecological dynamics, without pathogen evolution. Dynamics between seasons are represented, with infection starting at *t* = 4. In b), pathogen population is homogeneous, due to the absence of mutation. c) Eco-evolutionary dynamics with pathogen virulence evolution. Dynamics between seasons are represented, with infection starting at *t* = 4. Because of pathogen evolution, the impact of the infection in the last *h*_1_ provokes higher host density lost, compared to a). In d), time evolution of pathogen shows that the relative abundance of fitter strains – in darker colours – increases along seasons.

### 2.3 Sequence improvement: pattern rearrangement

Under eco-evolutionary dynamics, infection causes yield loss in susceptible cash crops, which becomes more noticeable when pathogen virulence increases. Consequently, pathogen presence has repercussions on each rotation sequence fitness: accumulation of cash is diminished, depending on the proportion of crops that survive the infection by the end of the season, making the fitness value decrease.

We need to compute how much yield is lost due to the pest, and for that, we have included a new parameter: the effective crop ratio, or *δ*_*i*_. It indicates the proportion of crops that contribute to the change in soil quality or cash yield. To represent the loss, we have set *δ*_*i*_ to be the ratio of the density of hosts remaining at the end of the season *h*_*i*_ and the initial host density *H*, Eq. (6),

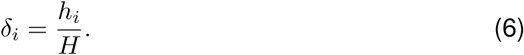

Included in the equations of soil quality and cash yield, Eqs. (7), *δ*_*i*_ modifies the outcome of the season.

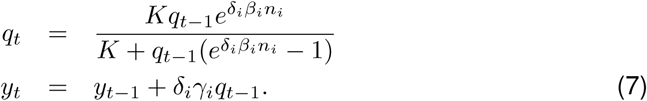

Therefore, in neutral conditions there is no loss of host density and *δ*_*i*_ = 1. In infection conditions, *δ*_1_ < 1 because of host death due to the pathogen; and *δ*_2_ = 1 as *h*_2_ density is not affected. The loss in host density is consequential for the final cash value, influencing the fitness *F* of each sequence.

Implementing eco-evolutionary dynamics in the model shows that under different infection scenarios (see Supplementary Material for infection at different time-points), the optimal sequence varies. Hence, the sequence with higher fitness under neutrality becomes non-optimal and can potentially be changed to a fitter sequence. Considering this, we next describe how to improve the initial sequence once infection has occurred.

Improving the current rotation sequence consists of comparing the focal optimal sequence – the best under neutral conditions – with the whole population under infection conditions. From the complete population of sequences, those ones which (i) patterns match until infection season – included –, and (ii) have greater fitness under infection, are considered candidate sequences for improvement. Since all infection scenarios show similarities among the optimal patterns (see Supplementary Material), few crop changes can already reduce the pest impact. As we assume no cost for pattern changes, we select the fittest sequence from the candidates (Fig. 5).

**Figure 5:**
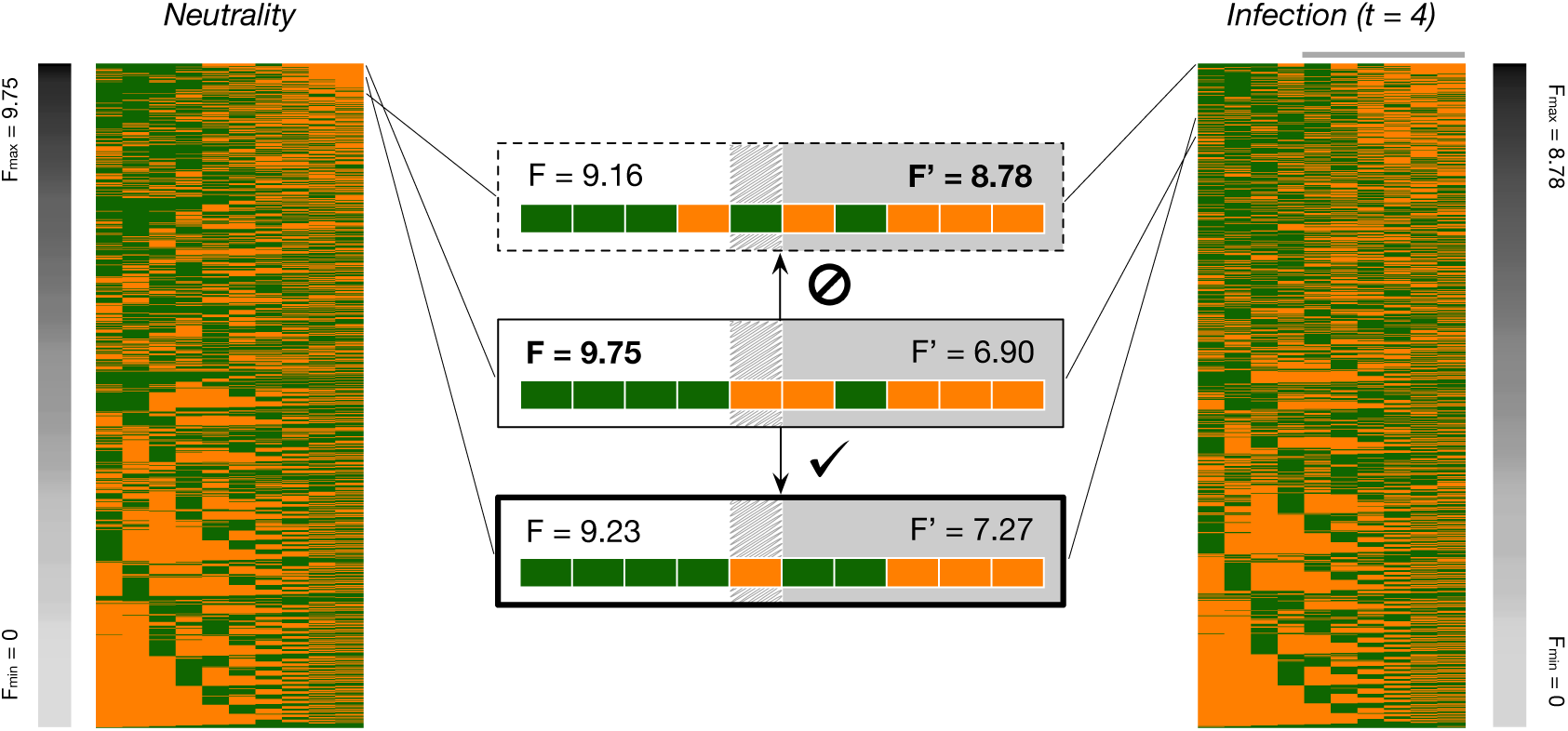
Pattern rearrangement strategies in infection conditions. In neutrality, the central sequence is calculated to be the fittest one (*F* = 9.75). However, when there is infection at *t* = 4 its fitness decays to *F* = 6.90. In infection conditions, the top sequence (dashed box) is indicated to be the fittest (*F* = 8.78); but because the pattern previous to the infection time (white background) is different, we cannot rearrange the pattern to be the best one. In consequence, bottom sequence (bold box) is the best option for minimising yield loss, as it is the fittest sequence (*F* = 7.27) where the pattern until the harvest of the infection crop is the same.

### 2.4 Sequence improvement: introduction of resistant cash crop

To prevent yield loss, a different strategy could be the introduction of a variant of the cash crop with resistance to the pathogen: 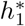. Acquiring a new resistance is associated with a cost, often a yield penalty [29, 30]. Thus, crop yield contribution would be set to 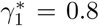 (in contrast to *γ*_1_ = 1, for the standard cash crop). However, the pathogen, in turn, could evolve to infect the resistant variant. The pathogen infecting multiple host types follows a gene-for-gene infection model [10].

Gene-for-gene dynamics has been widely described in plant-pathogen relationships [10, 31]. It is based on the idea that there is a correspondence between the genes providing resistance to the host and virulence for the pathogen. Carrying resistance supposes a cost for the host, and equivalently the pathogen has a cost for virulence. When acquiring the virulence allele, the pathogen does not lose its ability to infect the non-resistant host and thus, becomes a generalist. The contrasting model would be the matching alleles, in which the mutated pathogen is not infective any more for the host which genotype does not correspond specifically [18].

In our model, we modify the transition 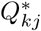 and fitness 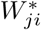 matrices to include gene-for-gene evolution of the pathogen. Each specialist pathogen *p*_*j*_ can mutate, with a transition probability *µ*, into its corresponding generalist variant 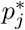 Furthermore, each strain of *p*_*j*_ and 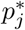 can mutate into fitter strains of their type (Fig. 6). In 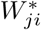, parameter *α* corresponds to the gradient between matching alleles and genefor-gene infection models, as described in [18]. We also include a cost *κ* of carrying a resistant allele for the mutated pathogen.

Replacing the previously susceptible cash crops *h*_1_ with resistant cash crops 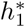, the frequency of newly evolved pathogens *p*_*j*_*\** increases until it outcompetes the old strains of pathogen *p*_*j*_ (Fig. 6c). Because of the cost of resistance of the evolved pathogen and the initial cash host resistance, the effect on yield loss is diminished. However, infection persists due to the adaptation of pathogen.

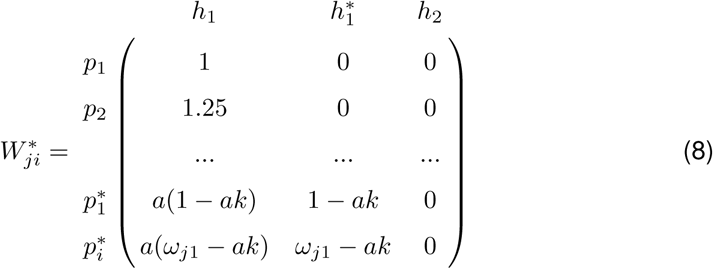

**Figure 6:**
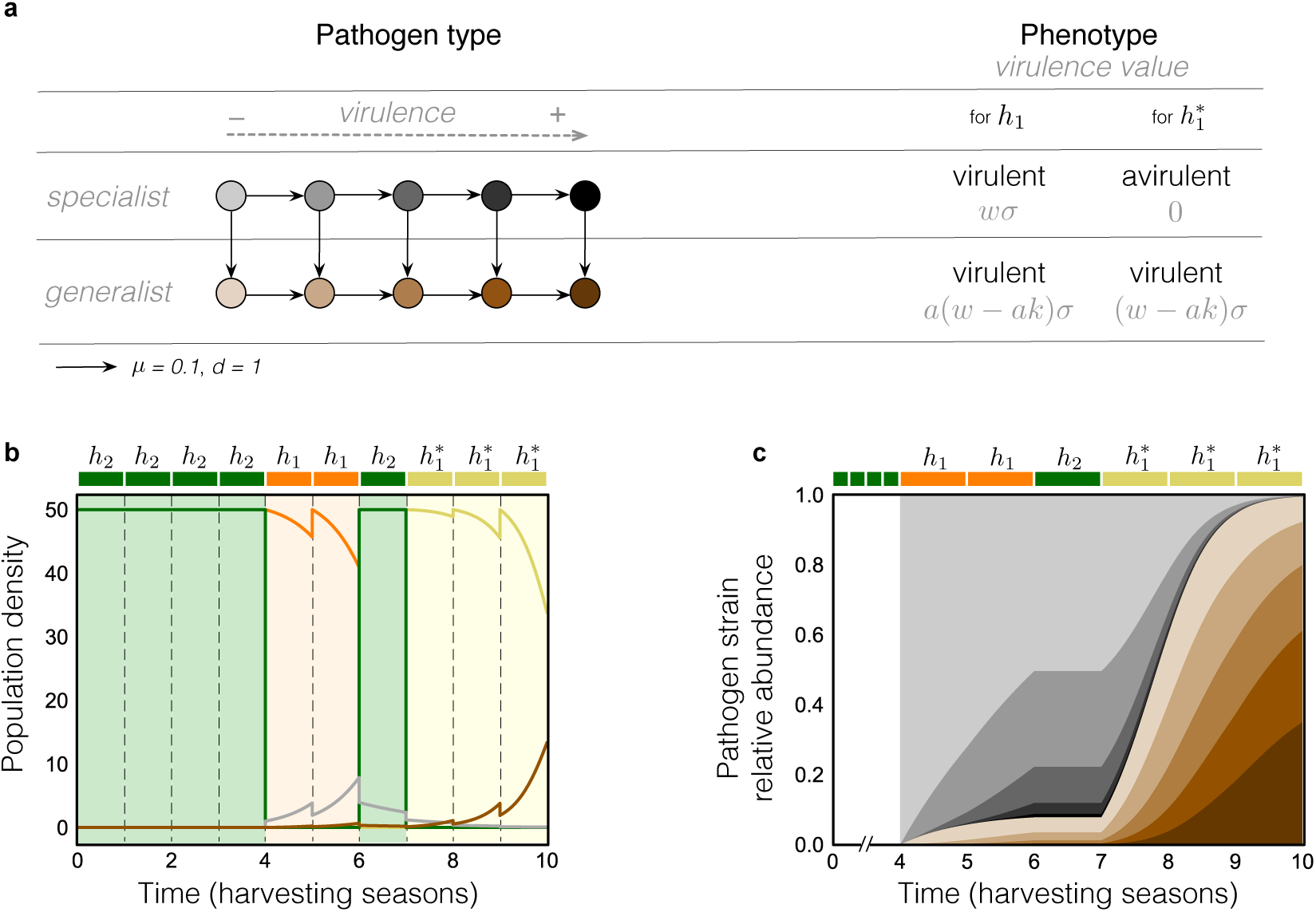
Pathogen gene-for-gene evolution and eco-evolutionary dynamics of host-pathogen interaction. a) Table for pathogen evolution, following gene-for-gene model of infection. b) Eco-evolutionary dynamics with pathogen virulence evolution, and adaptation to new host 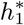. Dynamics between seasons are represented, with infection starting at *t* = 4. At *t* = 7, the new resistant host 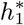 is cultivated, preventing the increase of pathogen density. However, the pathogen mutates following the gene-for-gene model, being able to infect both *h*_1_ and 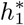, and at *t* = 8, there is a noticeable loss of host density. In c) strains of the original pathogen which only evolve in virulence – in grey – dominate pathogen population during the first seasons, in which only *h*_1_ and *h*_2_ are present. At *t* = 7, 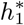 is introduced: only the strains which acquire the generalist mutation – in brown – are able to reproduce, which causes a fast increase on their relative abundance until the fixation of the generalist trait in the last season.

## Discussion

Translational evolutionary biology is an emerging field where fundamental concepts from evolutionary biology can be used in an applied setting to make practical changes in society [32, 33]. Just as with the search for novel antibiotics, the search for novel agricultural strategies has caught the interest of evolutionary biologists. Particularly, it has been discussed how evolutionary principles can be applied to pest management in agroecosystems. Our work complements previous attempts on coupling plant genetics with resistance deployment strategies [16, 34], bringing into focus plantpathogen dynamics and pathogen evolution in the context of crop rotation sequences.

We present a model for assessing how different patterns of cash and cover crop rotations influence long-term yield outcome. Our results highlight only a few patterns, from all possible crop sequences, maximising the yield after ten seasons of harvest. These patterns consist, overall, of investing in soil quality during the first seasons by cultivating cover crops, and once the carrying capacity of the soil is reached, introducing cash crops. For an accurate interpretation of these results, one must regard the conditions we have chosen. The initial soil quality, for example, is set at half of the carrying capacity, considering it an average value: different values could change the number of initial seasons invested in cover crops. Besides, using cover crops more efficient at increasing soil quality – this is, increasing their *β*_*i*_ – would increase the proportion of cash crops to be cultivated, augmenting the final cash yield. On the other hand, we have decided to maximise cash yield at the end of the sequence, for which last seasons are always dedicated to cash crops and soil quality is depleted. To allow continued use of the same sequence, we could optimise both soil quality and cash yield, so the final and initial soil quality have the same value. This new constraint would lead to a type of pattern possibly more regular in time, but reduced cash yield.

During the harvesting seasons, pathogens may invade the field and damage the crops, diminishing the expected yield. Using plant-pathogen dynamics, we have tracked the ecology and evolution of the infection in discrete and continuous time, predicting yield loss. Reported computational tools [7, 8, 9] rely on historical data to predict which is the best decision; instead, our model gives *de novo* assessment integrating features of the crop field, economy and evolution (i.e. soil quality, cash yield and pathogen dynamics, respectively). Moreover, we bridge evolutionary biology knowledge applied to agricultural strategies, whereas previous theoretical models are mostly focused on epidemiology [16, 35] and do not model explicitly pathogen dynamics and crop rotations sequences.

Farmers, looking for deployment strategies, can be guided by our improvement proposals: (i) pattern rearrangement and (ii) introduction of a resistant host. The first works purely at the ecological scale: it shows that by few changes in the optimal sequence based on reordering or adding cover crops, host resistance attenuates the increase of the pathogen load. This strategy can diminish the total yield loss, even when the number of cash crop seasons is reduced. The second approach integrates eco-evolutionary knowledge making use of resistant varieties of the crops arranged in the right pattern. It shows that using a new variant of cash crop that has resistance can help to overcome the pest during a certain time period. However, in the long term, the pathogen can adapt and infect the new crop type as well. We have implemented the co-evolutionary process following the gene-for-gene model of plantpathogen interactions [10]. Previous models describe the gene-for-gene approach in continuous evolutionary processes, whereas here it is used in the hybrid dynamical system across multiple harvesting seasons.

The agricultural strategies we propose fit previous work affirming that crop seasonality can slow down adaptation and promote evolutionary branching [36]. However, these strategies do not eradicate the infection, as in agriculture most of the crop pathogens evolve rapidly. This fast adaptation phenomenon occurs because high planting density and genetic uniformity of host increase the effective population size and genetic diversity of the pathogen [37]. Current research works on cultivar mixtures and their spatial distribution to diversify host genetics [38, 39, 40]. Our model considers a monoculture in a single field per season, but to complement these research lines, it could be extended by including more variation in host types, spatial structure, and between-field pathogen migration. Also, an increase in the number of host types and the number of pathogens could lead to a model exhibiting complex, and even chaotic, dynamics [41], which would be interesting to investigate.

Overall, our model can advise on strategies for maximising the gain of yield in cash crops, while using cover crops for soil improvement and control of pathogen spread. Further insights on rational resistance patterns could lead to new approaches for reducing pesticide use: the application of pesticides on strategic points where the pathogen density is low could both improve the efficiency of pesticides by increasing pest clearance and reducing the used amount of pesticide. These could possibly help disentangle the effect of crop rotations and pesticide use, in controlling pests and increasing yield. Experimental settings focus on crop rotations as the main factor for pest control, when put together with resistance variants and pesticides [22]. Agroecosystems rely on human artificial selection for controlling the outcome of the harvest: we can profit from an evolutionary outlook bringing new tools towards a more sustainable farming. Ideas such as the one presented in this research are merely the first steps towards achieving this goal.

## Code availability

Appropriate `Python` computer code describing the model is available at https://github.com/tecoevo/agriculture.

## Acknowledgements.

M.B-R and C.S.G. thank Alice Feurtey, Alexey Mikaberdize, Eva H. Stukenbrock and the department of Evolutionary Theory for fruitful discussions. Funding from the Max Planck Society is gratefully acknowledged.

## Supplementary material

### 4.1 *β* **parameter space**

The beta parameter indicates how fast a particular crop increases or decreases, according to its sign, the soil quality. Depending on its value, we expect the optimal patterns to be different: if cover crops improve soil quality rapidly, and cash crops decrease it very slowly, we would expect that optimal patterns have few cover crops, compared to the number of cash crops. In our case of study, we set *β*_1_ = *−*1.5 for cash crops (*h*_1_) and *β*_2_ = 1 for cover crops (*h*_2_), making soil quality decrease faster when one cash crop is cultivated, than the increase cover crops bring during one season. Using these *β* values, the optimal sequences presented a ratio of cash:cover crops of 1:1.

Here, we analyse the *β* parameter space in case of (i) the ratio of cash cover crops in the selection – sequences with fitness above two standard deviations over the global mean –, and (ii) the maximum fitness values of this selection. We do both analyses in the neutral scenario (without infection), and we explore the ratios coming from the combinations of values *β*_1_ = {−0.5, −1, −1.5, −2}, *β*_2_ = {0.5, 1, 1.5, 2}. The average values of the selection for each combination of *β* values is shown in a heatmap (Fig. SI.1).

Low *β*_1_ and high *β*_2_ lead to a higher proportion of cash crops in the rotation sequence and a general increase of fitness values. This can be interpreted as cover crops being very efficient, so few of them are needed to achieve values close to the carrying capacity of soil. Consequently, as fitness depends on the maximisation of cash, an increase on the proportion of cash crops – if the soil quality is high – brings to higher fitness. Combinations in which the absolute value of *β*_1_ equals *β*_2_, the ratio cash:cover is approximatively 1, leading to patterns and fitness maximum similar to the ones studied in the paper. Those cases in which *β*_1_ is greater than *β*_2_ tend to have low fitness maximum and low ratio cash:cover. Because of cover crops improving the soil quality slowly, many of them are needed to take profit of cultivating cash crops, reducing the cash season and thus, the economic outcome, or fitness.

### 4.2 Optimal sequence pattern depending on the time of infection

In sections 2.2 and 2.4, we have analysed eco-evolutionary dynamics showing a specific time of infection (*t* = 4). However, infection could occur at any season or time, and this could change the optimal sequence patterns. Here, we analyse the differences between the optimal patterns at different times of infection. Similarly to other sections, we have taken a subset of the population of sequences: those which fitness value *F* is over two standard deviations from the mean of the population. For each selection, we have plotted the frequencies of cash crops in each season (Fig. SI.2). During seasons in which frequency of cash equals 0 it is optimal to cultivate cover; per contra, if the frequency of cash is 1, it is always optimal to cultivate cash. When the frequency of cash is 0.5, the convenience of cash or cover depends on the crops cultivated before. However, we can take those cases as seasons in which choosing a particular type of crop will not change drastically the final outcome of the rotation.

**Figure SI.1:**
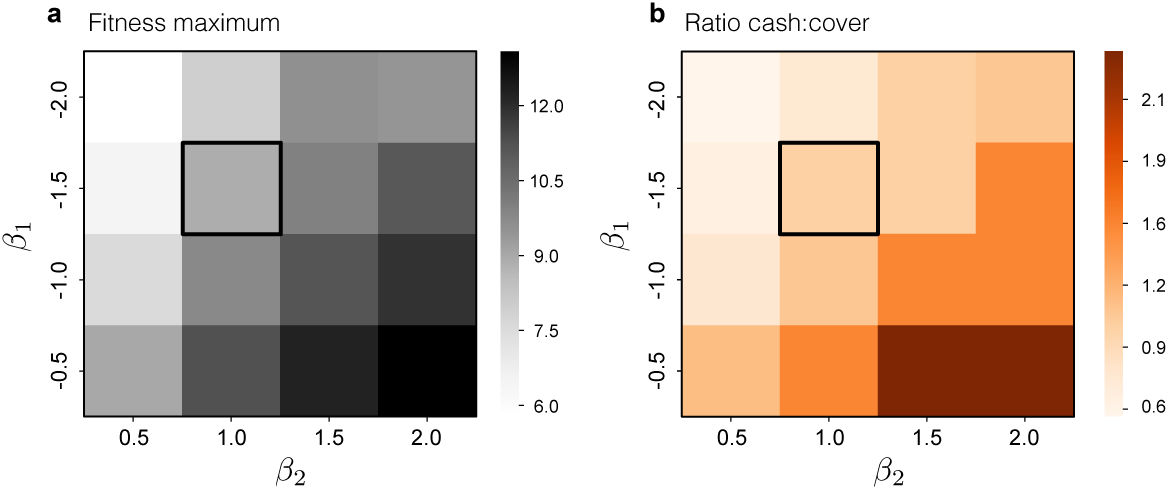
Fitness maximum and ratio cash:cover for different values of *β*_1_ and *β*_2_. Each patch in the heat map represents the mean value of (a) fitness maximum and (b) ratio cash:cover in the selection of optimal sequences – with fitness value higher than two standard deviations above the mean – in different combinations of *β*_*i*_ values, represented on the axis. The combination of values used for the results in the main text (*β*_1_ = *-*1.5, *β*_2_ = 1) is highlighted with a black square.

Results qualitatively show that the patterns are similar between them. In all of them, there is a tendency to start with cover crops and invest in cash crops at the end of the season; whereas the intermediate positions have intermediate frequency values and are variable among the optimal selection. Interestingly, optimal sequences do put a resistant cover crop after the time of infection in most of the cases, so pathogen load can be reduced. This is not the case for infection in the later times (*t* = 8, 9), in which keeping on investing in cash crops is yet the best strategy.

**Figure SI.2:**
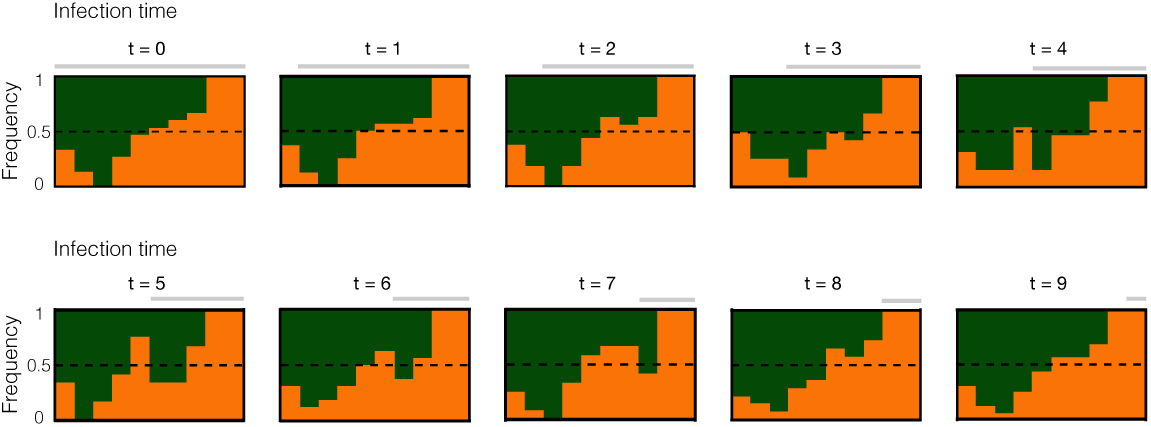
Frequency of cash crops in the optimal sequences at different times of infection. For each discrete time – i.e. season – of infection, the frequency of cash crops on the selection of optimal sequences (sequences with *F* over two standard deviations from the mean) is shown (orange bars). By subtracting the frequency of cash from 1, the frequency of cover crops can be inferred (corresponding to green in the plot). A frequency of 0 indicates that in all optimal sequences, cover crops are cultivated for that season. A frequency of 1 indicates that in all optimal sequences, cash crops are cultivated for that season. A frequency of 0.5 (dashed line) indicates that cover and cash crops are equally cultivated during that season. Other intermediate frequency values indicate that among optimal sequences, there is variation on the crop cultivated.

From this, we can observe that few changes in the optimal sequence in the neutral scenario can lead to optimal sequences in infection, and diminish fitness loss. Thus, the strategy of improvement based on pattern rearrangement is doable for all times of infection.

### 4.3 *α* parameter space

The *α* parameter refers to the gradient between gene-for-gene (GfG) and matching alleles (MA) models of infection. It was used in Agrawal and Lively (2002) [18] and posterior literature [23] for analysing the differences in behaviour of their systems between the two models of infection. Plant-pathogen coevolution is commonly associated to GfG patterns [10], and for our study we have used *α* = 0.5, which represents partial GfG. Here, we analyse how the pathogen strain relative abundance changes according the values *α* = {1, 0.5, 0}, corresponding to complete MA, intermediate GfG and complete GfG, respectively.

We compare the relative abundance of the pathogen *p*_1_ which can only infect *h*_1_ crops, and 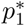 which can infect both *h*_1_ and 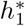. The analysis shows that the *α* value influences pathogen evolution. For *α* = 0, the pathogen stays specialist – following the MA model – until the 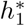 crop is introduced; then a mutated strain 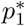 outcompetes the whole initial *p*_1_ population. No mutation nor abundance of 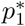 occurs before. For *α* = 0.5 we find an intermediate case: in the presence of *h*_1_ there is already low frequency (< 0.1) of *p*_1_*\**, which increases when 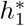 is introduced. For *α* = 1, frequency of *p*_1_*\** increases up to 0.2 before 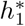 is introduced. With the introduction of 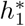, frequency of 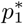 increases and reaches 1.

### 4.4 Fixed parameters

In the model, there are several parameters which have fixed values. We have compiled them in Table 1.

**Table 1:** List of fixed parameters used in the model.

| parameter | description | default value | reference |
| --- | --- | --- | --- |
| $q_0$ | Initial soil quality | $q_0 = 1$ | |
| $y_0$ | Initial cash yield | $y_0 = 0$ | |
| $K$ | Carrying capacity of the soil | $K = 2$ | |
| $\beta_i$ | Soil contribution of host $h_i$ | $\beta_1 = -1.5, \beta_2 = 1, \beta_1^* = -1.5$ | |
| $\gamma_i$ | Cash contribution of host $h_i$ | $\gamma_1 = 1, \gamma_2 = 0, \gamma_1^* = 0.9$ | |
| $\epsilon$ | Pathogen clearance | $\epsilon = 0.5$ | |
| $\kappa$ | Cost of resistance for the pathogen | $\kappa = 0.1$ | [18, 23] |
| $\mu$ | Transition probability | $\mu = 0.1$ | |
| $\alpha$ | GfG - MA gradient value | $\alpha = 0.5$ | [18, 23] |
| $\sigma$ | Infectivity | $\sigma = 0.04$ | [18, 23] |
| $H$ | Initial host density | $H = 50$ | |
| $d_p$ | Death rate of the pathogen | $d_p = 0.5$ | [18, 23] |

**Figure SI.3:**
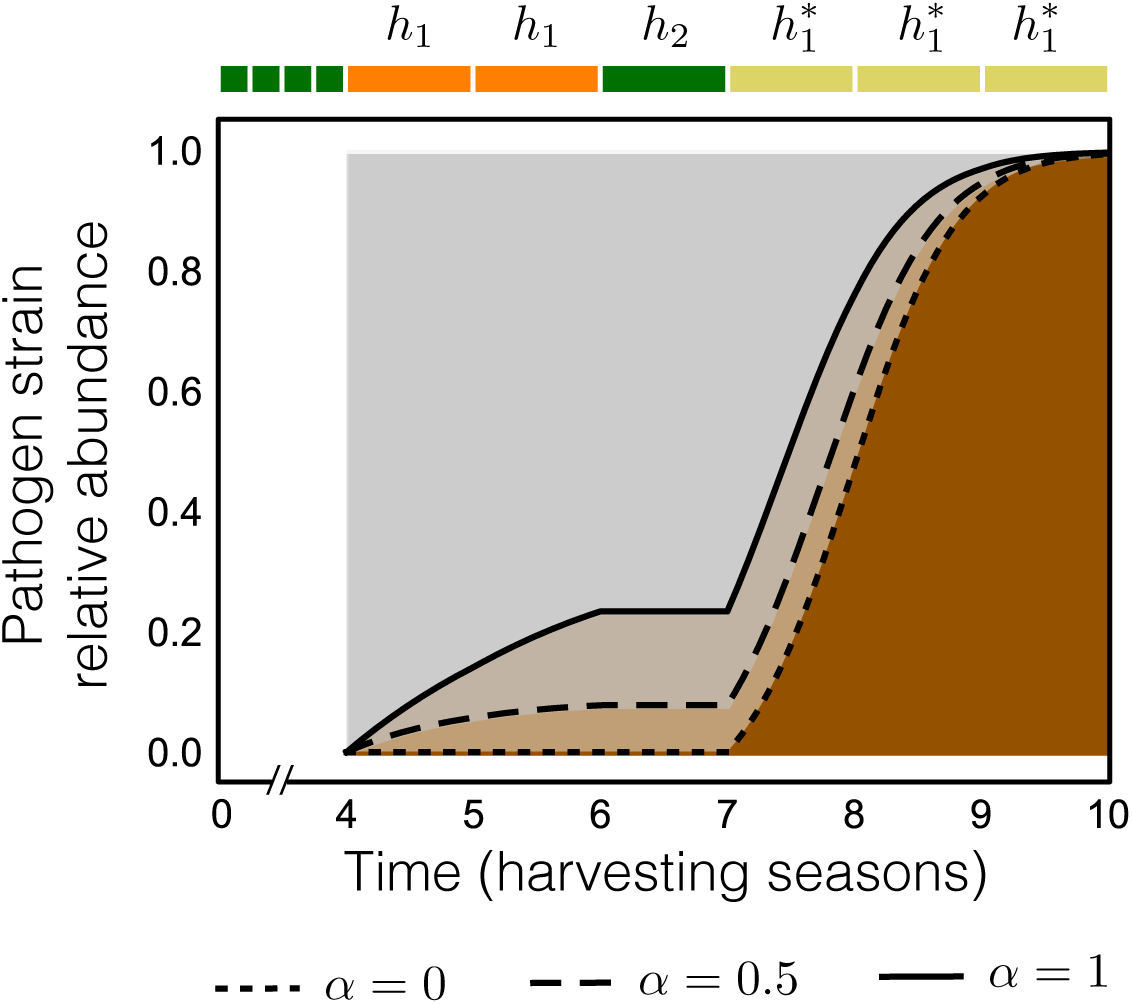
Pathogen evolution in different *α* values. Pathogen strain relative abundance is shown for three values of *α* (*α* = 0, 0.5, 1). When *α* = 0, the curve corresponds to a matching alleles (MA) model, in which the 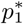 strain – in brown – only evolves when 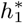 is present because it is specialised. On the contrary, *α* = 1 reflects a pure gene for gene (GfG) model, where the 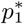 strain is a generalist and thus, already appears during the presence of *h*_1_. Value of *α* = 0.5 corresponds to an intermediate MA-GfG, and it is the value used in the main results.

## References

[1] Evans, L. T. Feeding the Ten Billion: Plants and Population Growth (Cambridge University Press, 1999).

[2] Oerke, E. C. Crop losses to pests. Journal of Agricultural Science. 144, 31–43 (2006).

[3] Nickel, J. L. Pest situation in changing agricultural systems a review. Bulletin of the Entomological Society of America 19, 136–142 (1973).

[4] Mohler, C. L. & Johnson, S. E. Crop rotation on organic farms: a planning manual., chap. How Expert Organic Farmers Manage Crop Rotations, 3–5 (Plant and Life Sciences Publishing, 2009).

[5] Harrison, P. World agriculture: towards 2015/2030, summary report. Tech. Rep., Food and Agriculture Organisation (2002).

[6] Foley, J. A. et al. Solutions for a cultivated planet. Nature 478, 337–342 (2011).

[7] Castellazzi, M. S. et al. A systematic representation of crop rotations. Agricultural Systems 97, 26–33 (2008).

[8] Osman, J., Inglada, J. & Dejoux, J. F. Assessment of a markov logic model of crop rotations for early crop mapping. Computers and Electronics in Agriculture 113, 234–243 (2015).

[9] Dury, J., Schaller, N., Garcia, F., Reynaud, A. & Bergez, J. E. Models to support cropping plan and crop rotation decisions. a review. Agronomy for Sustainable Development 32, 567–580 (2012).

[10] Flor, H. H. The complementary genetic systems in flax and flax rust. Advances in Genetics 8, 29–54 (1956).

[11] Grosberg, R. K. & Hart, M. W. Mate selection and the evolution of highly polymorphic self/nonself recognition genes. Science 289, 2111–2114 (2000).

[12] Thrall, P. H. et al. Evolution in agriculture: the application of evolutionary approaches to the management of biotic interactions in agro-ecosystems. Evolutionary Applications 4, 200–215 (2011).

[13] Neve, P., Vila-Aiub, M. & Roux, F. Evolutionary-thinking in agricultural weed management. New Phytologist 184, 783–793 (2009).

[14] Brown, J. K. M. & Tellier, A. Plant-parasite coevolution: Bridging the gap between genetics and ecology. Annual Review of Phytopathology 49, 345–367 (2011).

[15] Zhan, J., Thrall, P. H. & Burdon, J. J. Achieving sustainable plant disease management through evolutionary principles. Trends in Plant Science 19, 570–575 (2014).

[16] Burdon, J. J., Barret, L. G., Rebetzke, G. & Thrall, P. H. Guiding deployment of resistance in cereals using evolutionary principles. Evolutionary Applications 7, 609–624 (2014).

[17] Orr, H. A. & Unckless, R. L. Population extinction and the genetics of adaptation. The American Naturalist 172, 160–169 (2008).

[18] Agrawal, A. & Lively, C. M. Infection genetics: gene-for-gene versus matchingalleles models and all points in between. Evolutionary Ecology Research 4, 79–90 (2002).

[19] de Boer, H. C., van Eekeren, N. J. M., Pinxterhuis, J. B. & Stienezen, M. W. J. Optimal length of the grass-clover period in crop rotations: results of a 9-year field experiment under organic conditions. Tech. Rep. 660, Wageningen UR Livestock Research (2012).

[20] Day, T. & Otto, S. P. Fitness. In Encyclopedia of Life Sciences (Nature Publishing Group, 2001).

[21] Anderson, P. K. et al. Emerging infectious diseases of plants: Emerging infectious diseases of plants: pathogen pollution, climate change and agrotechnology drivers. Trends in Ecology and Evolution 19, 535–544 (2004).

[22] Marburger, D. A. et al. Crop rotation and management effect on fusarium spp. populations. Crop Science 55, 365–376 (2015).

[23] Song, Y., Gokhale, C. S., Papkou, A., Schulenburg, H. & Traulsen, A. Hostparasite coevolution in populations of constant and variable size. BMC Evolutionary Biology 15, 212 (2015).

[24] van der Schaft, A. J. & Schumacher, H. An Introduction to Hybrid Dynamical Systems, vol. 251 of Lecture Notes in Control and Information Sciences (Springer, 2000).

[25] Carroll, S. P., Hendry, A. P., Resnick, D. N. & Fox, C. W. Evolution on ecological time-scales. Functional Ecology 21, 387–393 (2007).

[26] Pelletier, F., Garant, D. & Hendry, A. P. Eco-evolutionary dynamics. Philosophical Transactions of the Royal Society B: Biological Sciences 364, 1483–1489 (2009).

[27] Frickel, J., Sieber, M. & Becks, L. Eco-evolutionary dynamics in a coevolving host–virus system. Ecology Letters 19, 450–459 (2016).

[28] Gandon, S., van Baalen, M. & Jansen, V. A. A. The evolution of parasite virulence, superinfection, and host resistance the evolution of parasite virulence, superinfection, and host resistance the evolution of parasite virulence, superinfection, and host resistance. The American Naturalist 159, 658–669 (2002).

[29] Brown, J. K. M. Yield penalties of disease resistance in crops. Current Opinion in Plant Biology 5, 339–344 (2002).

[30] Burdon, J. J. & Thrall, P. H. The fitness costs to plants of resistance to pathogens. Genome Biology 4, 227 (2003).

[31] Thompson, J. N. & Burdon, J. J. Gene-for-gene coevolution between plants and parasites. Nature 360, 121–125 (1992).

[32] Fang, F. C. & Casadevall, A. Lost in translation–basic science in the era of translational research. Infection and immunity 78, 563–566 (2010).

[33] Carroll, S. P. et al. Applying evolutionary biology to address global challenges. Science 346, 1245993 (2014).

[34] Zhan, J., Thrall, P. H., Papaix, J., Xie, L. & Burdon, J. J. Playing on a pathogen’s weakness: Using evolution to guide sustainable plant disease control strategies. Annual Review of Phytopathology 53, 19–43 (2015).

[35] Papaix, J., Burdon, J. J., Zhan, J. & Thrall, P. H. Crop pathogen emergence and evolution in agro-ecological landscapes. Evolutionary Applications 8, 385–402 (2015).

[36] Hamelin, F. M., Castel, M., Poggi, S., Andrivon, D. & Mailleret, L. Seasonality and the evolutionary divergence of plant parasites. Ecology 92, 2159–2166 (2011).

[37] McDonald, B. A. & Stukenbrock, E. H. Rapid emergence of pathogens in agroecosystems: global threats to agricultural sustainability and food security. Philosophical Transactions of the Royal Society B: Biological Sciences 371, 20160026 (2016).

[38] Mikaberidze, A., McDonald, B. A. & Bonhoeffer, S. Developing smarter host mixtures to control plant disease. Plant Pathology 64, 996–1004 (2014).

[39] Fabre, F., Rousseau, E., Mailleret, L. & Moury, B. Epidemiological and evolutionary management of plant resistance: optimizing the deployment of cultivar mixtures in time and space in agricultural landscapes. Evolutionary Applications 8, 919–32 (2015).

[40] Djidjou-Demasse, R., Moury, B. & Fabre, F. Mosaics often outperform pyramids: Insights from a model comparing strategies for the deployment of plant resistance genes against viruses in agricultural landscapes. New Phytologist 216, 239–253 (2017).

[41] Schenk, H., Traulsen, A. & Gokhale, C. S. Chaotic provinces in the kingdom of the Red Queen. Journal of Theoretical Biology 431, 1–10 (2017).

